# Exploring the relationships between autozygosity, educational attainment, and cognitive ability in a contemporary, trans-ancestral American sample

**DOI:** 10.1101/2021.11.24.469902

**Authors:** Sarah MC Colbert, Matthew C Keller, Arpana Agrawal, Emma C Johnson

**Affiliations:** Department of Psychiatry, Washington University School of Medicine, Saint Louis, MO; Department of Psychology, University of Colorado Boulder, Boulder, CO; Institute for Behavioral Genetics, University of Colorado Boulder, Boulder, CO

**Keywords:** runs of homozygosity, autozygosity, educational attainment, assortative mating, cognitive ability

## Abstract

Previous studies have found significant associations between estimated autozygosity - the proportion of an individual’s genome contained in homozygous segments due to distant inbreeding - and multiple traits, including educational attainment (EA) and cognitive ability. In one study, estimated autozygosity showed a stronger association with parental EA than the subject’s own EA. This was likely driven by parental EA’s association with mobility: more educated parents tended to migrate further from their hometown, therefore choosing more genetically diverse partners. We examined the associations between estimated autozygosity, cognitive ability, and parental EA in a contemporary sub-sample of adolescents from the Adolescent Brain and Cognitive Development Study^SM^ (ABCD Study^®^) (analytic N=6,504). We found a negative association between autozygosity and child cognitive ability consistent with previous studies, while the associations between autozygosity and parental EA were in the expected direction of effect (with greater levels of autozygosity being associated with lower EA) but the effect sizes were significantly weaker than those estimated in previous work. We also found a lower mean level of autozygosity in the ABCD sample compared to previous autozygosity studies, which may reflect overall decreasing levels of autozygosity over generations. Variation in migration and mobility patterns in the ABCD study compared to other studies may explain the pattern of associations between estimated autozygosity, EA, and cognitive ability in the current study.

## Introduction

Runs of homozygosity (ROHs) are stretches of DNA that are identical by descent (Wright, 1922); these arise when an individual’s parents share a distant common ancestor. While greater numbers of and longer ROHs tend to be associated with more recent inbreeding (e.g., cousin-cousin inbreeding), ROHs occur even in seemingly outbred populations (McQuillan et al., 2008). Previous studies have found that individuals with a greater level of autozygosity (F_ROH_; the proportion of the genome contained in ROHs) tend to have lower values on fitness-related traits, such as cognitive ability (Howrigan et al., 2016; Joshi et al., 2015), respiratory function (e.g., forced expiratory volume; Johnson et al., 2018; Joshi et al., 2015), and reproductive characteristics such as number of offspring (Clark et al., 2019; Johnson et al., 2018; Yengo et al., 2017). This phenomenon, known as inbreeding depression, suggests that these traits have been under selection pressures over evolutionary time, biasing the genetic variants that influence those traits towards being rare and recessive.

ROHs have been used to examine evolutionary hypotheses about quantitative traits (e.g., cognitive performance) and case-control phenotypes (e.g., psychiatric diagnoses). For example, multiple studies have shown that increased autozygosity is associated with decreased cognitive ability (Abdellaoui et al., 2015; Howrigan et al., 2016), consistent with the hypothesis that genetic variants that negatively impact cognitive ability have been under directional selection pressures and are more likely to be rare and recessive, and thus exert their effects in regions of homozygosity. Associations have been less consistent for case-control phenotypes such as schizophrenia (Johnson et al., 2016; Keller et al., 2012), potentially due to ascertainment differences in cases and controls and/or sociodemographic factors that may play a role in assortative mating (Abdellaoui et al., 2013; Clark et al., 2019). To adequately account for these potential confounding factors requires familial data—either parent-offspring or sibling (Clark et al., 2019; Johnson et al., 2018)—but this type of familial phenotypic and genotypic data is rarely available in large enough samples to achieve adequate statistical power.

Abdellaoui et al. demonstrated the utility of including both parental and offspring phenotypes in ROH analyses in a 2015 study where they found a stronger negative association between the offspring’s autozygosity and *parental* educational attainment (EA) (p < 9e-5) than between the offspring’s autozygosity and their own EA (p = 0.045). This negative association between parental EA and offspring autozygosity was entirely mediated by the distance between parental birthplaces, with evidence to suggest that more highly educated parents tended to be more mobile on average and thus less likely to mate with an individual who shares a distant common ancestor, potentially inducing F_ROH_ ~ trait associations that are due to sociological factors rather than reflecting a true effect of inbreeding depression.

The Adolescent Brain Cognitive Development Study^SM^ (ABCD Study^®^) is a large, longitudinal study with genetic and phenotypic data available for approximately 11,000 adolescents, as well as limited phenotypic data on their parents. The baseline sample includes children who were 9-11 years old in 2016-2017, and their parents. As such, this sample provides us an opportunity to examine (1) the distribution of autozygosity in a contemporary North American sample and (2) whether increased autozygosity is associated with decreased cognitive ability and parental EA.

## Methods

### Preregistration

We preregistered our analysis (osf.io/eqkty). Data and analyses presented in this manuscript closely resemble those described in the preregistration; any exceptions are described here. First, while we perform analyses separately for genetically confirmed non-Hispanic European ancestry and non-Hispanic African ancestry samples, we eventually chose to meta-analyze across ancestry groups. Deviations in sample sizes reflect a better understanding of the information available for each individual as we began implementing the analyses in the data. We also recoded the parental EA measure using codes different from those in the preregistration so that our measure of EA might more closely resemble that used by Abdellaoui et al. (2015). Lastly, we included additional sensitivity analyses not described in the preregistration which are strictly exploratory and can be found in the Supplementary Note.

### Sample

This study used genetic and phenotypic data from the ABCD study version 3.0, a long-term study of brain development through adolescence in over 11,000 children (Jernigan et al., 2018). Data were initially collected in children ages 9-10 across 21 sites and have subsequently been collected each year. Genetic samples were collected from the children in addition to demographic and phenotypic data on both the children and their parents.

Using a combination of self-report race and ethnicity and genetic principal component analysis (PCA), we separated the 11,875 ABCD Study participants into a predominantly European genetic ancestry subset (N = 5,556) and predominantly African genetic ancestry subset (N = 1,584), to account for differences in allele frequencies and linkage disequilibrium across populations (see *Genotypic Data Cleaning*), which can lead to differences in mean F_ROH_ across ancestry groups. We then called runs of homozygosity (ROHs) and estimated F_ROH_ (the proportion of the genome contained in ROHs) for the PCA-selected European- and African-ancestry individuals separately (see *ROH Calling*). To maximize sample size in each analysis, we included any individual if data for the necessary variables (outcome and covariates) were available. For example, if EA data were available for an individual’s mother but not father, that individual was included in the test for an association between child’s F_ROH_ and maternal EA, but excluded from the sample used to test for an association between child’s F_ROH_ and paternal EA. As such, Ns varied depending on data availability for each individual and specific Ns are defined for each analysis in Table S1.

### Phenotypes

Parental EA was measured using the question “What is the highest grade or level of school you have completed or the highest degree you have received?” This field was only considered for individuals for whom we could confirm that the answer was for the biological parent of the child. To approximate the coding for EA in Abdellaoui et al., responses were recoded into the three categories: (1) Completed High school, GED or less: included individuals who reported never attending kindergarten or that they completed a grade between first and 12th, High school, a GED or equivalent diploma; (2) Completed higher education up to Bachelor’s Degree: included individuals who reported completing some college, an occupational Associate degree, an academic Associate degree or a Bachelor’s degree; and (3) Completed Master’s or Doctoral degree: included individuals who reported completing a Master’s or Doctoral degree.

Given the age of the children (current ages ranging from 14-15) in the study, the above definitions of EA were not applicable to the ABCD Study child subjects. Although EA is a multi-faceted construct, there is substantial evidence for a strong correlation between educational achievement and cognitive ability (Deary et al., 2007; Kaufman et al., 2009; Lynn & Meisenberg, 2010; Strenze, 2007), and previous studies have identified significant associations between F_ROH_ and cognitive ability (Clark et al., 2019; Howrigan et al., 2016; Johnson et al., 2018; Yengo et al., 2017); therefore, we also examined the association between F_ROH_ and the child’s cognitive ability, measured by their Overall Cognition Composite Score (Akshoomoff et al., 2013; Weintraub et al., 2013). We chose to use the score uncorrected for age, as age was already included as a covariate in our model (see *Statistical Analysis*).

### Genotypic data cleaning

We used the Rapid Imputation and COmputational PIpeLIne for Genome-Wide Association Studies (RICOPILI; Lam et al., 2020) to perform quality control (QC) on the 11,099 individuals with available ABCD Study phase 3 genotypic data, using RICOPILI’s default parameters. The 10,585 individuals who passed QC checks were then matched to broad self-report racial groups using the ABCD Study parent survey. There were 6,787 individuals for whom their parents/caregivers indicated the child’s race was only “white”, and 5,561 of those individuals did not endorse any Hispanic ethnicity/origin. We also identified 1,675 individuals for whom their parents/caregivers indicated the child’s race was only “black”, and 1,584 of those individuals did not endorse any Hispanic ethnicity/origin. After performing a second round of QC on these sub-samples, 5,556 non-Hispanic White and 1,584 non-Hispanic Black individuals were retained in the analyses. Principal component analysis (PCA) in RICOPILI (Lam et al., 2020) was used to confirm the genetic ancestry of these individuals by mapping onto the 1000 Genomes reference panel (Auton et al., 2015), resulting in PCA-selected European- and African-ancestry subsets.

### Statistical analysis

Statistical analyses consisting of ROH calling, F_ROH_ estimation and association testing were performed separately for the PCA-selected European- and African-ancestry subsets. Results of association tests were then meta-analyzed across the two ancestry groups using a fixed-effect model implemented with the “metafor” package in R (Viechtbauer, 2010). We first tested for heterogeneity using a random-effects model for each meta-analysis; however, tests for heterogeneity were non-significant (p>0.05), thus we chose to use fixed-effect models over random-effect models. We report the meta-analysis results as the main findings.

#### ROH calling

Following the procedures of previous studies (Clark et al., 2019), we cleaned the data further using PLINK 1.9 (Chang et al., 2015), excluding SNPs with > 3% missingness or a MAF < 5 % and excluding individuals with > 3% missing data. After QC, 288,246 SNPs remained for the EUR sample and 278,639 SNPs remained for the AFR sample.

We called ROHs using PLINK 1.9 (Chang et al., 2015), following the approach taken in Abdellaoui et al. (2015) and the recommendations of Howrigan et al. (2011). We first pruned SNPs for LD (window size = 50, number of SNPs to shift after each step = 5, based on a variance inflation factor [VIF] of 2) using the following parameters in PLINK 1.9: --indep 50 5 2. After LD pruning, 93,952 and 136,164 SNPs were left for analysis of the EUR and AFR samples, respectively. Next, we defined an ROH as ≥ 65 consecutive homozygous SNPs, with no heterozygote calls allowed using the following PLINK code: --homozyg-window-het 0 -- homozyg-snp 65 --homozyg-gap 500 --homozyg-density 200.

We also called ROHs using the method presented by Clark et al. (2019), which differs not only in the ROH calling procedure, but also in that there is no initial LD pruning. To call ROHs we used the following parameters in PLINK 1.9: --homozyg-window-snp 50; --homozyg-snp 50; --homozyg-kb 1500; --homozyg-gap 1000; --homozyg-density 50; --homozyg-window-missing 5; homozyg-window-het 1. Results using this method are presented in the Supplementary Note.

#### F_ROH_ calculation and association analysis

F_ROH_ was calculated as the total length of ROHs summed for each individual, and then divided by the total SNP-mappable autosomal distance (2.77 × 10^6^ kilobases). We used mixed effect regression models to test the association between (1) child’s F_ROH_ and child’s cognitive ability and (2) child’s F_ROH_ and parental EA (both maternal and paternal EA, separately). In each linear regression model, child’s F_ROH_ was the outcome variable and cognitive ability, maternal EA or paternal EA was included as a predictor variable. To account for the non-normal distribution of F_ROH_, we calculated empirical p-values using a permutation procedure (as in Abdellaoui et al.) implemented in the permlmer package (Lee & Braun, 2012) in R and ran 10,000 permutations to calculate an empirical p-value for each model. All empirical p-values were nearly identical to observed p-values in the original models; thus, we meta-analyzed the original effect sizes. Each model included the following covariates: child’s age, child’s biological sex, genotyping batch, testing site, family ID, and the first ten ancestry PCs. Testing site and family ID were modeled as random intercepts and all other covariates were fixed. We also performed a test for association between child’s F_ROH_ and child’s cognitive ability while accounting for maternal and paternal EA as additional covariates.

## Results

### Descriptive statistics

We called ROHs and estimated F_ROH_ in 5,556 PCA-selected European-ancestry individuals and 1,584 PCA-selected African-ancestry individuals (7,140 individuals total). The extent of inbreeding in the ABCD sample overall was quite low, with an average F_ROH_ of 0.00052 (SD = 0.00378), minimum F_ROH_ of 0 (6,081 individuals had zero ROHs) and maximum F_ROH_ of 0.077. We also computed the number of ROH segments, with the number of ROHs in each individual ranging from 0 to 26 and averaging at 0.234. The average total amount of ROH — that is, the combined length of all ROH segments — was 1.43 MB. Ancestry-specific estimates are available in the Supplementary Note. For descriptive statistics from child cognition and parental EA please see Table S2.

### Autozygosity, educational attainment, and cognitive ability in the ABCD Study sample

#### Association between child cognitive ability and child F_ROH_

Of the 7,140 individuals with ROH calls, F_ROH_ estimates, and information on covariates, data on child cognitive ability were available for 6,504 individuals. Child cognitive ability was negatively associated with F_ROH_ in the primary meta-analysis across ancestry groups (standardized beta = −0.032, standard error = 0.014, p = 0.022). Results were similar when inbreeding outliers (F_ROH_ > 0.0156) were excluded, with the meta-analysis showing a negative association between child cognitive ability and F_ROH_ (standardized beta = −0.032, standard error = 0.013, p = 0.014). In general, both the PCA-selected European- and African-ancestry subsamples showed consistent direction of effect, but standard errors were larger in the much smaller PCA-selected African-ancestry subset (all results provided in Table 1). Using the ROH calling method from Clark et al. (2019), we also detected a negative association between child cognitive ability and F_ROH_ (see Supplementary Note). We also tested the association between child cognitive ability and F_ROH_ while accounting for both maternal and paternal EA in 3,983 individuals who had data for all three phenotypes, and found that the effect size was identical to the analysis where we did not control for parental EA, although the standard error increased (standardized beta = −0.032, standard error = 0.017, p = 0.058).

#### Associations between parental educational attainment and child F_ROH_

Of the 7,140 individuals with ROH calls, F_ROH_ estimates, and information on covariates, data on maternal EA and paternal EA were available for 5,801 and 4,172 individuals, respectively. In the primary meta-analysis, maternal EA was negatively associated with F_ROH_, although the association was not statistically significant (standardized beta = −0.02, standard error = 0.013, p = 0.120; after removing inbreeding outliers: beta = −0.012, standard error = 0.014, p = 0.402). Similarly, paternal EA was negatively associated with F_ROH_, but this association was not statistically significant (standardized beta = −0.029, standard error = 0.017, p = 0.082; after removing inbreeding outliers: standardized beta = −0.007, standard error = 0.017, p = 0.679).

## Discussion

In a sample of approximately 7,000 adolescents of PCA-selected European- and African ancestries, we found a negative association between child F_ROH_ and child cognitive ability (standardized beta = −0.032; s.e. = 0.014; p = 0.022), replicating previous findings (Abdellaoui et al., 2015; Howrigan et al., 2016); however, we note that this p-value would not withstand Bonferroni correction for 3 tests (association tests between F_ROH_ and cognitive scores, paternal EA, and maternal EA; α = 0.0167). Effect sizes were slightly smaller but of similar magnitude in our study (standardized beta = −0.032; s.e. = 0.014) relative to Abdellaoui et al.’s study (standardized beta = −0.041; s.e. = 0.024). Notably, the negative association we identified between child F_ROH_ and child cognitive ability was of similar magnitude when maternal and paternal EA were included as covariates in the model.

While several previous studies suggest that a negative association between cognitive ability and F_ROH_ may indicate inbreeding depression on cognitive ability (Howrigan et al., 2016), Abdellaoui et al. suggested that their findings were the result of individuals with lower EA being less likely to migrate and more likely to mate with others who are also less educated and less mobile, leading to parents having more similar genetic backgrounds on average and their child therefore displaying more autozygosity while also being genetically predisposed to lower educational attainment. In support of this hypothesis, they found significant associations between child F_ROH_ and both maternal EA (standardized beta = −0.080; s.e. = 0.021) and paternal EA (standardized beta = −0.089; s.e. = 0.022) that were stronger than the association between F_ROH_ and the child’s own EA (standardized beta = −0.041; s.e. = 0.024). However, not only do we find that our parent EA-child F_ROH_ associations are significantly weaker (yet still in the same direction of effect) compared to the parent EA-child F_ROH_ associations found in Abdellaoui et al., but in our sample, child F_ROH_ is more strongly associated with child cognitive ability (standardized beta = −0.032; s.e. = 0.014) than either maternal EA (standardized beta = - 0.020; s.e. = 0.013) or paternal EA (standardized beta = −0.029; s.e. = 0.017).

While we do not rule out lack of power as a contributing factor (Keller et al., 2011), we deem it unlikely to be the sole explanation for our weaker results for parental EA measures, as the current sample size was large (N ~ 7,000) relative to Abdellaoui et al.’s study (N ~ 2,000), which found a highly significant association between F_ROH_ and both maternal and paternal EA. Given the smallest reported effect size in Abdellaoui et al.’s study (standardized beta = −0.041, s.e. = 0.024) and the reported N = 2,007, we estimate that we would have 84% power to detect an effect of the same size in our sample of 5,181 European ancestry individuals, given that the standard deviation of F_ROH_ is very similar across the two studies: 0.0031 in our European-ancestry subset vs. 0.003 in Abdellaoui et al. Below, we consider other possible explanations for our differing results.

Notably, the current sample and the sample used in Abdellaoui et al., 2015 were derived from two different countries (the United States and the Netherlands, respectively), which bear varying degrees of resemblance in terms of cultural, social, and economic contexts. The possible influence of these differences across countries are evident in previous studies which have found opposite directions of associations between autozygosity and various phenotypes. For example, previous research has identified a negative association between F_ROH_ and EA in a Dutch population (Abdellaoui et al., 2015) and cognitive ability in a broader European population (Howrigan et al., 2016), and oppositely, a positive association between F_ROH_ and cognitive ability in both an American sample (Córdova-Palomera et al., 2018) and a UK sample (Power et al., 2014). We consider that the results of our current study, which uses a contemporary American sample, may differ from previous findings partly as a result of the sample’s demographics.

An important aspect of the Abdellaoui et al. study is the discovery that migration is a significant mediator in the relationship between parental EA and child’s F_ROH_. Individuals with higher EA on average had traveled a greater distance between their birthplace and their spouse’s birthplace, as well as between their birthplace and their child’s birthplace (Abdellaoui et al., 2015). That is to say, EA and mobility were positively associated, resulting in individuals with higher EA being somewhat more likely to mate with more genetically dissimilar individuals on average. As a result, the offspring of individuals with higher EA may be more outbred as well as inheriting a predisposition for higher EA. In support of this theory, Abdellaoui et al. found that the association between parental EA and child’s F_ROH_ was fully mediated by the distance between maternal and paternal birthplace, although birthplaces for both offspring and parents only included locations within the Netherlands. While the Abdellaoui et al. study only considered internal migration, the ABCD sample includes children born to individuals who may have migrated internationally, not just within the same country, as parents in the ABCD study born in a wide variety of places, which range from the United States (data on exact location not provided) to countries like Mexico and Yemen. We did not have information available on city or state-specific places of birth, leaving us unable to investigate the relationships between EA, mobility, and F_ROH_ in our sample. While the impact of migration and its relationships with F_ROH_ and EA were potentially more straightforward and interpretable in the context of domestic migration in the Netherlands, complicated political and historical contexts which influence international migration and differ country to country likely produce variable relationships between EA and autozygosity, and the lack of state- or even region-specific data limit our ability to draw conclusions and comparisons in the ABCD data.

Incidentally, we found very low levels of autozygosity in the ABCD sample compared to previous studies. Whereas the variance of F_ROH_ was similar to that of previous studies, the mean F_ROH_ observed in the current study (F_ROH_ = 0.0005) was much lower than that observed in Abdellaoui et al. (F_ROH_ = 0.0016), Howrigan et al. (F_ROH_ = 0.0041) and Power et al. (F_ROH_ = 0.007). While the mean F_ROH_ in a sample is not immediately pertinent to the statistical power to detect associations between F_ROH_ and complex traits, we suspect that generational differences in mean F_ROH_ may still be relevant to the findings of this and other studies which examine F_ROH_. The participants of the ABCD study were primarily born between 2006 and 2007, with the median birth year of *parents* of ABCD individuals being 1976, compared to Abdellaoui et al.’s study in which two-thirds of the offspring studied were born in 1984 or earlier. To our knowledge, there are few ROH studies that have included young (birth year > 1990) United States samples, but one study of 809 North Americans of European descent aged 19-99 years old found that ROH significantly decreased in size and frequency as chronological age decreased; furthermore, the authors predicted a decline in percent F_ROH_ of ~0.1 for every 20 years’ difference in birth year (Nalls et al., 2009). Given that the ABCD participants are around 20 years younger than the subjects in Abdellaoui et al., and the average percent F_ROH_ in Abdellaoui et al.’s sample was ~0.16, we might expect the average percent F_ROH_ in ABCD to be ~0.06, or an average F_ROH_ of 0.0006; this is similar to the observed average F_ROH_ of 0.0005.

We conducted our own assessment of this phenomenon by comparing several generations within another North American sample (the Collaborative Study on the Genetics of Alcoholism (COGA) (Begleiter et al., 1995)) to assess differences in autozygosity over time. In the youngest generation, which included individuals born after 1990 (the youngest individual in this sample was born in 2003), the average F_ROH_ was 0.0002 (s.e. = 3.2e-5), while the average F_ROH_ calculated in older generations were 0.0003 (s.e. = 2.6e-5), 0.0007 (s.e. = 1.1e-4), 0.0008 (s.e. = 1.4e-4), 0.0010 (s.e. = 2.8e-4) and 0.0034 (s.e. = 3.4e-3) for those born between 1970-1990, 1950-1970, 1930-1950, 1910-1930 and 1890-1910, respectively (Table S3). We also ran a linear mixed model which further confirmed a significant effect of birth year on F_ROH_ (beta = - 0.071, s.e. = 0.011, p = 4.3e-11). Furthermore, we compared two cohorts from similar geographic regions in Howrigan et al.’s 2016 study to assess differences in autozygosity according to generation in a UK sample, to see if the pattern held across cultures. In the young (average age=11.67 years old) GAIN UK cohort, the average F_ROH_ was 0.0019, while the average F_ROH_ calculated in the older English MANC cohort (average age = 64.9 years old) was 0.0047 (from Tables 1 and 2 in Howrigan et al.).

Therefore, we hypothesize that year of birth may be impacting F_ROH_ estimates across studies; given the exponential increases in population size and urbanization (Bongaarts, 2009) as well as a decline in racial and religious endogamy (Kalmijn, 1991; Luo, 2017; Rosenfeld, 2008) both globally and in the US, all of these factors may be contributing to decreasing levels of inbreeding over time (Bittles & Black, 2010; Campbell et al., 2009; Nalls et al., 2009). Thus, the generational difference between our study sample, ABCD, and those of previous ROH studies may partly explain the lower levels of autozygosity observed in the current study. The ABCD sample was genotyped on the Smokescreen array (Baurley et al., 2016), which is built on an Affymetrix backbone with additional addiction-focused content. As Abdellaoui et al.’s sample was genotyped on the Affymetrix 6.0 array, and we followed identical genotyping QC and ROH calling procedures, it seems unlikely that differences in SNP panel or calling algorithms are responsible for the lower levels of autozygosity observed in our study.

In summary, we found a negative association between estimated autozygosity and child cognitive ability, such that individuals with lower estimates of F_ROH_ tended to have higher levels of cognitive ability, replicating previous findings. On the other hand, we found weaker associations between F_ROH_ and maternal EA or paternal EA, although findings were generally in the expected, negative direction of effect. We hypothesize that these mixed results are due to a combination of generational differences in autozygosity and the complex mechanisms which influence both EA and mobility in different countries. Future studies should carefully characterize and consider how the effects of assortative mating, migration patterns, and generational differences in the distribution of F_ROH_ may influence autozygosity-trait associations across samples.

## Acknowledgements

We thank Sarah Paul for her help with phenotype curation.

Data used in the preparation of this article were obtained from the Adolescent Brain Cognitive Development^SM^ (ABCD) Study (https://abcdstudy.org), held in the NIMH Data Archive (NDA). This is a multisite, longitudinal study designed to recruit more than 10,000 children age 9-10 and follow them over 10 years into early adulthood. The ABCD Study^®^ is supported by the National Institutes of Health and additional federal partners under award numbers U01DA041048, U01DA050989, U01DA051016, U01DA041022, U01DA051018, U01DA051037, U01DA050987, U01DA041174, U01DA041106, U01DA041117, U01DA041028, U01DA041134, U01DA050988, U01DA051039, U01DA041156, U01DA041025, U01DA041120, U01DA051038, U01DA041148, U01DA041093, U01DA041089, U24DA041123, U24DA041147. A full list of supporters is available at https://abcdstudy.org/federal-partners.html. A listing of participating sites and a complete listing of the study investigators can be found at https://abcdstudy.org/consortium_members/. ABCD consortium investigators designed and implemented the study and/or provided data but did not necessarily participate in the analysis or writing of this report. This manuscript reflects the views of the authors and may not reflect the opinions or views of the NIH or ABCD consortium investigators.

We also thank the Collaborative Study on the Genetics of Alcoholism for generously allowing us to include an analysis of their data in this manuscript. The Collaborative Study on the Genetics of Alcoholism (COGA), Principal Investigators B. Porjesz, V. Hesselbrock, T. Foroud; Scientific Director, A. Agrawal; Translational Director, D. Dick, includes eleven different centers: University of Connecticut (V. Hesselbrock); Indiana University (H.J. Edenberg, T. Foroud, Y. Liu, M. Plawecki); University of Iowa Carver College of Medicine (S. Kuperman, J. Kramer); SUNY Downstate Health Sciences University (B. Porjesz, J. Meyers, C. Kamarajan, A. Pandey); Washington University in St. Louis (L. Bierut, J. Rice, K. Bucholz, A. Agrawal); University of California at San Diego (M. Schuckit); Rutgers University (J. Tischfield, R. Hart, J. Salvatore); The Children’s Hospital of Philadelphia, University of Pennsylvania (L. Almasy); Virginia Commonwealth University (D. Dick); Icahn School of Medicine at Mount Sinai (A. Goate, P. Slesinger); and Howard University (D. Scott). Other COGA collaborators include: L. Bauer (University of Connecticut); J. Nurnberger Jr., L. Wetherill, X., Xuei, D. Lai, S. O’Connor, (Indiana University); G. Chan (University of Iowa; University of Connecticut); D.B. Chorlian, J. Zhang, P. Barr, S. Kinreich, G. Pandey (SUNY Downstate); N. Mullins (Icahn School of Medicine at Mount Sinai); A. Anokhin, S. Hartz, E. Johnson, V. McCutcheon, S. Saccone (Washington University); J. Moore, Z. Pang, S. Kuo (Rutgers University); A. Merikangas (The Children’s Hospital of Philadelphia and University of Pennsylvania); F. Aliev (Virginia Commonwealth University); H. Chin and A. Parsian are the NIAAA Staff Collaborators. We continue to be inspired by our memories of Henri Begleiter and Theodore Reich, founding PI and Co-PI of COGA, and also owe a debt of gratitude to other past organizers of COGA, including Ting-Kai Li, P. Michael Conneally, Raymond Crowe, and Wendy Reich, for their critical contributions. This national collaborative study is supported by NIH Grant U10AA008401 from the National Institute on Alcohol Abuse and Alcoholism (NIAAA) and the National Institute on Drug Abuse (NIDA).

## Declarations

### Ethics approval

This study was approved by the local Institutional Review Board.

### Consent to participate

All participants in ABCD provided informed consent (or assent).

### Consent for publication

Not applicable

### Availability of data and material

Genetic and phenotypic data in the ABCD sample are available for download for approved researchers from the NIMH Data Archive.

### Author contributions

Study design and conception were developed by Sarah MC Colbert and Emma C Johnson, with input from Matthew C Keller. Data cleaning and preparation and statistical analyses were performed by Sarah MC Colbert. Analyses were supervised by Arpana Agrawal and Emma C Johnson. All authors participated in critical discussion of the results. The manuscript draft was written, edited, and approved by all authors.

