## Supplementary material for "Exploring the relationships between autozygosity, educational attainment, and cognitive ability in a contemporary, trans-ancestral American sample": Table 1

**Table 1.** Association between child F_ROH_ and maternal EA, paternal EA and child’s cognitive ability. Associations were tested both before and after excluding individuals with hypothesized second-cousin inbreeding.

|  | ***Primary trans-ancestral meta-analysis*** | | ***PCA-selected European-ancestry*** | | ***PCA-selected African-Ancestry*** | |
| --- | --- | --- | --- | --- | --- | --- |
| **Model** | **Standardized Beta (SE)** | **P** | **Standardized Beta** | **P** | **Standardized Beta (SE)** | **P** |
| Child cognitive ability | -0.03 (0.01) | 0.022 | -0.04 (0.02) | 0.013 | -4.79e-03 (0.03) | 0.869 |
| Maternal EA | -0.02 (0.01) | 0.120 | -0.03 (0.01) | 0.027 | 0.03 (0.03) | 0.343 |
| Paternal EA | -0.03 (0.02) | 0.082 | -0.03 (0.02) | 0.043 | -0.02 (0.05) | 0.714 |
| Child cognitive ability with parent EA covariates | -0.03 (0.02) | 0.058 | -0.04 (0.02) | 0.039 | 0.07 (0.06) | 0.260 |
| ***Excluding hypothesized second-cousin inbreeding outliers (F_roh_ > 0.015625)*** | | | | | | |
| Child cognitive ability | -0.03 (0.01) | 0.014 | -0.03 (0.02) | 0.034 | -0.04 (0.03) | 0.145 |
| Maternal EA | -0.01 (0.01) | 0.402 | -9.43e-3 (0.02) | 0.540 | -0.02 (0.03) | 0.425 |
| Paternal EA | -0.007 (0.02) | 0.679 | -0.01 (0.02) | 0.433 | 0.02 (0.05) | 0.714 |
