## Supplementary Note for "Exploring the relationships between autozygosity, educational attainment, and cognitive ability in a contemporary, trans-ancestral American sample"

**Supplemental Note**

Ancestry specific Descriptive Statistics

*F_ROH_*

In the PCA-selected European ancestry sub-sample and African ancestry-sub sample the average F_ROH_ = 0.00038 (SD = 0.0031) and 0.00098 (SD = 0.0054), respectively. The average number of ROHs in the PCA-selected European ancestry sub-sample was 0.191 and 0.388 for the PCA-selected African ancestry-sub sample. The average total amount of ROH was 1.06 MB and 2.72 MB for the PCA-selected European and African ancestry sub-samples, respectively.

Replication of Analysis Using Clark et al. Method

*Descriptive statistics*

We called ROHs and estimated F_ROH_ in 5,556 PCA-selected European-ancestry individuals and 1,584 PCA-selected African-ancestry individuals (7,140 individuals total), this time using the method presented by Clark *et al.* (2019). In general, we observed slightly higher F_roh_ estimates using this method, with the average F_roh_ being 0.00078, minimum F_ROH_ of 0 (4,622 individuals had zero ROHs) and maximum F_ROH_ of 0.081. We also computed the number of ROH segments, with the number of ROHs in each individual ranging from 0 to 19 and averaging at 0.5756, over two times larger than the average that was observed in the main findings. Most likely to be contributing to these differences is the lower threshold for the length of homozygous SNPs required to define an ROH. The method used by Abdellaoui *et al.* (2015) and presented by Howrigan *et al.* (2011) defines an ROH as a segment with > 65 homozygous SNPs, whereas the Clark *et al.* (2019) method includes segments with > 50 homozygous SNPs.

*Association between child cognitive ability and child F_ROH_*

Using the Clark et al. method, child cognitive ability was negatively associated with F_ROH_ in the primary meta-analysis (beta = -7.9e-6, standard error = 3.5e-6, p = 0.023). Similar to the main findings, both the PCA-selected European- and African-ancestry subsamples showed consistent direction of effect, but standard errors were larger in the much smaller PCA-selected African-ancestry subset.

*Associations between parental educational attainment and child F_ROH_*

As was done using the ROH calls computed using the Howrigan *et al.* (2011) method, we tested for significant associations between parental EA and child F_ROH_. In the primary meta-analysis, maternal EA was not significantly associated with F_ROH_, (beta = -7.6e-5, standard error = 5.1e-5, p = 0.168); however, paternal EA was significantly associated with F_ROH_ (beta = -1.4e-5, standard error = 5.3e-5, p = 0.011). Once again, we observed that associations between parental EA measures and F_ROH_ were significant in the PCA-selected European-ancestry subsamples, but not in the PCA-selected African-ancestry subsamples.

Exploration of Age-by-F_ROH_ association in COGA

To explore whether the low average autozygosity we observed in the ABCD sample (relative to previous ROH studies) might be due to a hypothesized decrease in autozygosity over generations, we examined the distribution of F_ROH_ in an independent comparison sample, the Collaborative Study on the Genetics of Alcoholism (COGA). While there are no participants in COGA who are as young as those in ABCD, COGA is still a relatively ideal sample for this comparison: there is a wide range of birth years – from 1890 to 2003 – in COGA, and it is also a diverse sample drawn from sites across the United States, similar to ABCD. More detailed descriptions of COGA are available elsewhere (Begleiter *et al.*, 1995); briefly, the COGA sample was ascertained for probands with alcohol use disorders and their close relatives from seven US sites. Control families were recruited from the community. Institutional review boards from all seven sites approved the study and all participants provided informed consent. COGA samples were genotyped on four different SNP arrays, and the genotype data were QCed using standard procedures (see Lai *et al.*, 2019 for futher details). For the following analyses, we filtered to only include SNPs that were also available in the ABCD sample for consistency across studies.

First, we calculated the average F_ROH_ across generations (Table S2), observing an overall decrease in F_ROH_. Next, we formally tested this in a linear mixed model (controlling for sex, 10 genetic PCs and array type as fixed covariates, and family ID as a random intercept) and found a significant effect of birth year on F_ROH_ in both the European (beta = -0.085, s.e. = 0.014, p = 2.6e-9) and African ancestry subsets (beta = -0.051, s.e. = 0.017, p = 3.6e-3). We meta-analyzed these results to test the effect in the overall sample and present these results in the main manuscript (beta = -0.071, s.e. = 0.011, p = 4.3e-11). These findings provide additional evidence that there appears to be an overall decrease in autozygosity over generations, at least in samples from the U.S.

**Table S1.** Sample descriptions. Number of individuals available for each variable, or groups of variables (e.g., there are 4,259 genotyped individuals for which we have data on cognitive ability and each of the covariates included in our model).

| **Variable(s)** | | **N PCA-selected European-ancestry** | **N PCA-selected African-ancestry** | **N total** |
| --- | --- | --- | --- | --- |
| Genetic Data | | 5,556 | 1,584 | 7,140 |
|  | Child Cognitive Ability | 5,181 | 1,323 | 6,504 |
|  | Maternal EA | 4,623 | 1,178 | 5,801 |
|  | Paternal EA | 3,770 | 402 | 4,172 |

**Table S2A.** Descriptive statistics for child cognitive ability.

| **Variable** | **Mean** | **Minimum** | **Maximum** |
| --- | --- | --- | --- |
| Child cognitive ability | 86.99 | 44 | 117 |

**Table S2B.** Descriptive statistics for parental educational attainment measures. Asterisks indicate levels of educational attainment for which the maternal and paternal portions significantly differed.

| **Level of Educational Attainment** | **N_maternal_** | **Proportion_maternal_** | **N_paternal_** | **Proportion_paternal_** |
| --- | --- | --- | --- | --- |
| High school, GED, or less* | 686 | 0.12 | 522 | 0.13 |
| Higher education up to Bachelor’s degree* | 3,622 | 0.62 | 2,497 | 0.60 |
| Completing a Master’s or Doctoral degree | 1,493 | 0.27 | 1,123 | 0.29 |

**Table S3.** Average F_ROH_ across generations in the Collaborative Study on the Genetics of Alcoholism.

| **Birth year** | **N** | **Average F_ROH_ (SE)** | **European ancestry subset Average F_ROH_ (SE)** | **African ancestry subset Average F_ROH_ (SE)** |
| --- | --- | --- | --- | --- |
| 1890 - 1910 | 4 | 0.00339 (3.39e-3) | 0.00339 (3.39e-3) | N/A |
| 1910 - 1930 | 236 | 0.00104 (2.83e-4) | 0.00117 (4.74e-3) | 0.00046 (2.27e-4) |
| 1930 - 1950 | 897 | 0.00077 (1.42e-4) | 0.00087 (4.87e-3) | 0.00049 (1.15e-4) |
| 1950 - 1970 | 2806 | 0.00074 (1.12e-4) | 0.00057 (4.30e-3) | 0.00093 (2.04e-4) |
| 1970 - 1990 | 2663 | 0.00031 (2.64e-5) | 0.00025 (1.45e-3) | 0.00038 (3.60e-5) |
| 1990+ | 1347 | 0.00023 (3.18e-5) | 0.00022 (1.38e-3) | 0.00024 (3.32e-5) |
